# Characterization of the urinary DNA virome of hematopoietic stem cell transplant recipient and healthy cynomolgus macaques

**DOI:** 10.64898/2026.05.05.722665

**Authors:** Helena Vogel, Tristan T Neal, Helen L Wu, Kaitlyn Kukula, Paul Kievit, Jonah Sacha, Gabriel J Starrett

**Affiliations:** Laboratory of Cellular Oncology, CCR, NCI, NIH, Bethesda, Maryland, USA; Vaccine and Gene Therapy Institute, Oregon Health & Science University, Beaverton, Oregon, USA; Oregon National Primate Research Center, Oregon Health & Science University, Beaverton, Oregon, USA

## Abstract

BK polyomavirus (BKPyV) and chronic latent DNA virus infections frequently reactivate after hematopoietic stem cell transplantation (HSCT), where they contribute to major complications in humans with limited options for prevention or treatment. Mauritian cynomolgus macaques (MCM; *Macaca fascicularis*) serve as an important animal model for transplantation research due to their limited major histocompatibility complex (MHC) diversity. Similar viral reactivations and disease complications occur in MCMs and humans post-transplantation, including polyomavirus-associated hemorrhagic cystitis and nephropathy. To define the polyomaviruses associated with these outcomes and characterize the broader urinary DNA virome, we sequenced rolling circle-amplified urine DNA from HSCT recipient and healthy cynomolgus macaques, allowing for comprehensive genomic characterization of the detected polyomaviruses. De novo assembly and annotation identified three polyomaviruses with apparent specific host tropism to cynomolgus macaques: *Macaca fascicularis* polyomavirus 2 (MafaPyV2) and *Macaca fascicularis* polyomavirus 3 (MafaPyV3), which are closely related to human polyomaviruses, and a newly identified strain of simian virus 40 (SV40 type IIB). All three viruses were detected in both HSCT recipients and healthy animals but were shed at significantly higher relative loads in HSCT recipients. Multiple polyomaviruses were frequently detected within individual hosts, and all three were identified in HSCT recipients that developed urologic disease. Together, these findings further characterize polyomavirus diversity, shedding, and disease associations in hematopoietic stem cell–transplanted Mauritian cynomolgus macaques.

**Importance:** Reactivation of latent viruses after hematopoietic stem cell transplantation is a major cause of illness in transplant recipients, yet options for prevention and treatment remain limited. Progress has been hindered by the lack of animal models that accurately reflect human disease. In this study, we show that Mauritian cynomolgus macaques harbor multiple polyomaviruses that reactivate at high levels following transplantation and are associated with urinary tract disease, closely paralleling complications seen in humans. We define the genetic identity of these viruses and demonstrate that immunosuppression strongly enhances viral shedding and disease severity. By establishing clear similarities between viral reactivation in macaques and humans, this work validates the Mauritian cynomolgus macaque transplantation model as a powerful platform for studying virus-driven disease mechanisms and testing antiviral therapies and vaccines relevant to human transplantation.

## Introduction

In humans, many DNA viruses increase in abundance during states of immune suppression, such as in the context of HIV/AIDS, cancer therapy, and solid organ or hematopoietic stem cell transplantation (HSCT). Viral reactivation in these settings can lead to severe disease, such as nephropathy and hemorrhagic cystitis, driven by uncontrolled replication of normally subclinical infections of polyomaviruses, adenoviruses, or herpesviruses^1^. Specifically, BK polyomavirus (BKPyV), a ubiquitous DNA virus that establishes a lifelong infection in the urinary tract of humans, can replicate to high viral titers after transplantation and lead to hemorrhagic cystitis and/or BKPyV-associated nephropathy (BKVN), characterized primarily by renal dysfunction or hematuria and histologic evidence of viral cytopathic injury^1,2^. Clinical disease often requires immunosuppression reduction and can result in graft rejection and exacerbation of graft-versus host disease (GVHD)^3^. Despite the clinical impact of these infections, therapeutic and preventative interventions are extremely limited and opportunistic viral infections remain a major cause of morbidity and mortality after solid organ or HSCT^4^. Nonhuman primates provide essential models for transplantation research because they closely recapitulate human immune and physiological responses. Mauritian cynomolgus macaques (MCM; *Macaca fascicularis*) represent a particularly valuable model due to their limited genetic diversity, especially at major histocompatibility complex (MHC) loci, resulting from a small founder population introduced to Mauritius more than 500 years ago^5–7^. An allogeneic HSCT model developed in MCMs has previously been shown to mirror key features of human post-transplant infectious complications, including viral reactivation and urologic disease caused by a macaque polyomavirus analogous to human BKPyV^8^. In this model, some HSCT recipients develop hematuria, hemorrhagic cystitis, and renal dysfunction, with occasional histologic evidence of viral cytopathic changes and immunoreactivity for polyomavirus large T antigen in the kidney^8^. The virus detected in these animals has historically been referred to as “cynomolgus polyomavirus” (CPV), first described in 1999 in immunosuppressed cynomolgus macaques with interstitial nephritis following renal transplantation^9^. However, these prior studies relied on PCR and Sanger sequencing of short, conserved regions of the large T antigen gene, leaving the complete genomic composition, diversity, and number of polyomaviruses circulating in HSCT recipient and healthy cynomolgus macaques unresolved. As a result, the genetic relationships among these viruses, their host specificity, and their potential association with disease remain incompletely defined.

In this study, we aimed to comprehensively characterize the DNA virome and the diversity and genomic features of polyomaviruses shed by HSCT recipient and healthy MCMs. Using rolling circle amplification followed by short-read Illumina and long-read Nanopore sequencing, we performed a retrospective analysis of urine samples from HSCT recipients and healthy controls, as well as plasma and spleen samples from one HSCT recipient with severe hemorrhagic cystitis and polyomavirus-associated nephritis. Through de novo genome assembly and annotation, we resolved the complete genomes of multiple cynomolgus macaque polyomaviruses, characterized their diversity and phylogenomics, and assessed their prevalence and relative abundance in immunocompromised and immunocompetent hosts.

The close parallels between viral reactivation and disease manifestations in MCMs and human transplant recipients establish this HSCT model as a powerful platform for studying polyomavirus biology, viral evolution under immune suppression, and mechanisms of polyomavirus-associated disease. Furthermore, comprehensive genomic characterization of these viruses provides a foundation for improving diagnostic approaches and for evaluating preventive and therapeutic interventions, including vaccines and antiviral strategies, in the setting of transplantation.

## Results

### Urinary DNA virome of HSCT recipient and healthy cynomolgus macaques

De novo assembly and annotation of rolling circle amplified DNA sequencing data from 80 temporally discrete urine samples collected from 16 HSCT recipient MCMs and 42 healthy cynomolgus macaques (34 Mauritian, 6 Philippine, 1 Cambodian, and 1 Indonesian-origin macaque) revealed marked differences in urinary virome composition between groups (Table 1). The cumulative urinary virome from HSCT recipients was dominated by polyomaviruses, followed by bacteriophages, phylogenetically unclassified viruses, CRESS DNA viruses, and anelloviruses (Fig. 1A-C). At lower relative abundances, papillomavirus, parvovirus, herpesvirus, varidnavirus, caudoviricetes, and mastadenovirus were detected (Table S2). Cumulatively, healthy control animals predominantly shed bacteriophage, unclassified viruses, and CRESS DNA viruses (Fig. 1A, D) with polyomavirus, anellovirus, papillomavirus, herpesvirus, varidnavirus and hepadnavirus detected at low relative abundance (Table S2). Within the HSCT recipient samples, the predominance of polyomavirus reads was primarily driven by five samples from two animals (CM67 and CM69) (Fig. 1B). A few samples from healthy animals were shedding polyomavirus, however, the average relative abundance was only 320 polyomavirus reads per million total reads (VRPM), compared to an average of 137,000 VRPM in HSCT recipients (Fig. 1B-D and Table S1). When excluding CM67 and CM69, the average relative abundance of polyomavirus shedding in HSCT-recipients remains higher than controls, at 5,359 VRPM. Considering the previously identified polyomavirus in HSCT recipient MCMs with urinary disease through Sanger sequencing and renal large T antigen immunoreactivity, subsequent analysis focused on comprehensive genomic characterization of detected polyomaviruses. Three distinct polyomaviruses were identified in HSCT recipient MCMs and healthy controls and named *Macaca fascicularis* polyomavirus 2 (MafaPyV2), *Macaca fascicularis* polyomavirus 3 (MafaPyV3), and SV40 type IIB. Alignment of complete genomes generated in this study with previously published short Sanger sequences of cynomolgus polyomavirus (CPV)^8^ revealed that these past sequences actually represented two different polyomaviruses, MafaPyV2 and SV40 type IIB (Fig. S1).

**Fig. 1.**
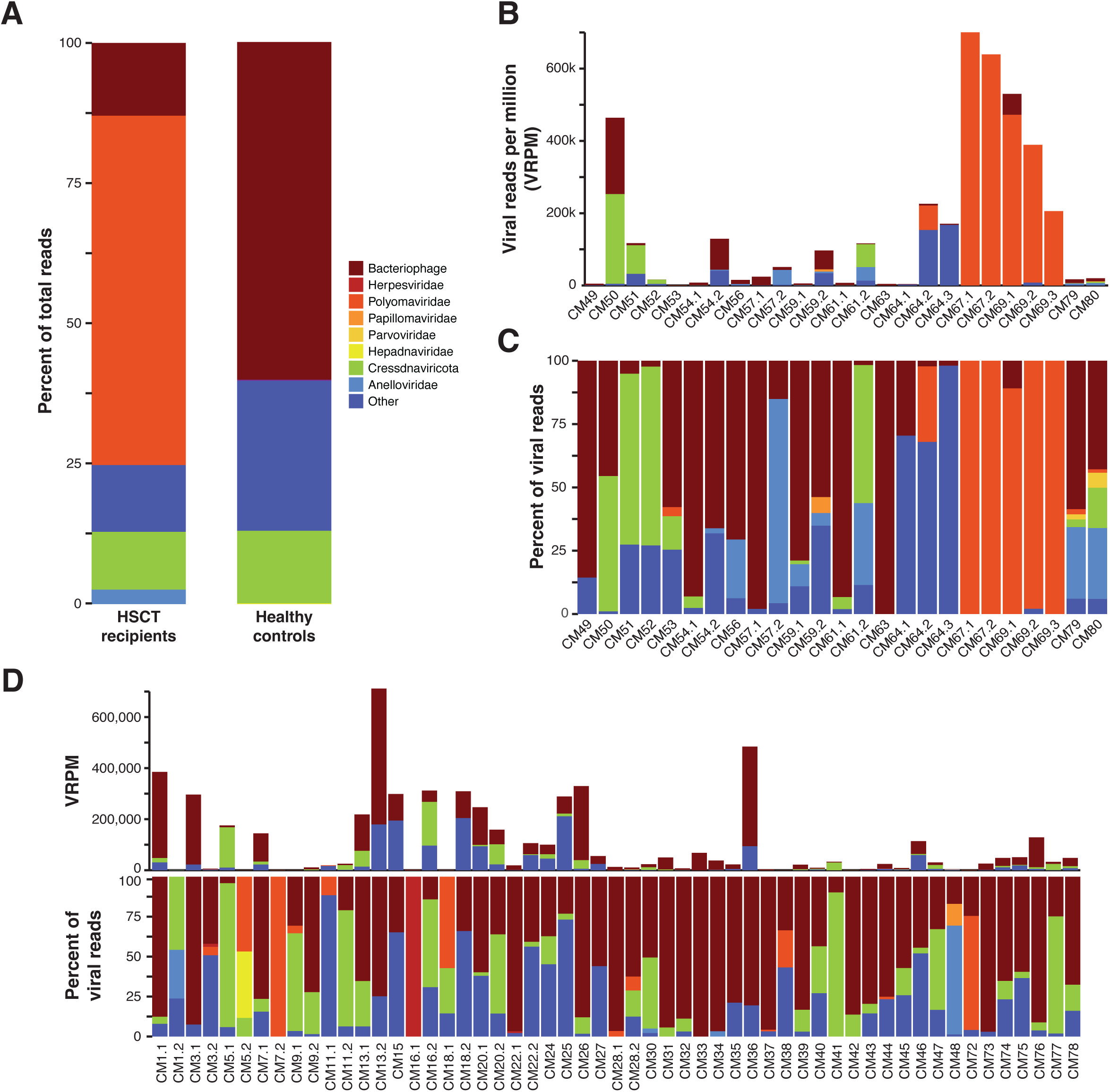
Viral group abundance in urine. (A) Viral family abundance in urine from HSCT recipient and healthy cynomolgus macaques including viruses present in >1% of total reads. Viral groups with <1% abundance are listed in Table S2. (B) Abundance of viral groups shed in urine of HSCT-recipients represented by viral reads per million (VRPM). (C) Percentage of reads in each viral group shed in urine of HSCT-recipients. (D) Viral family abundance in urine from healthy control cynomolgus macaques in VRPM and as percent of reads.

**Table 1.** Cynomolgus macaque sample collection data.

| Sample ID | Animal ID | Geographic origin | Sample type | HSCT-recipient or control |
| --- | --- | --- | --- | --- |
| CM1 | CM1.1 | Indonesian | Urine | control |
| CM2 | CM1.2 | Indonesian | Urine | control |
| CM3 | CM3.1 | Cambodian | Urine | control |
| CM4 | CM3.2 | Cambodian | Urine | control |
| CM5 | CM5.1 | Mauritian | Urine | control |
| CM6 | CM5.2 | Mauritian | Urine | control |
| CM7 | CM7.1 | Mauritian | Urine | control |
| CM8 | CM7.2 | Mauritian | Urine | control |
| CM9 | CM9.1 | Mauritian | Urine | control |
| CM10 | CM9.2 | Mauritian | Urine | control |
| CM11 | CM11.1 | Mauritian | Urine | control |
| CM12 | CM11.2 | Mauritian | Urine | control |
| CM13 | CM13.1 | Mauritian | Urine | control |
| CM14 | CM13.2 | Mauritian | Urine | control |
| CM15 | CM15 | Mauritian | Urine | control |
| CM16 | CM16.1 | Mauritian | Urine | control |
| CM17 | CM16.2 | Mauritian | Urine | control |
| CM18 | CM18.1 | Mauritian | Urine | control |
| CM19 | CM18.2 | Mauritian | Urine | control |
| CM20 | CM20.1 | Mauritian | Urine | control |
| CM21 | CM20.2 | Mauritian | Urine | control |
| CM22 | CM22.1 | Mauritian | Urine | control |
| CM23 | CM22.2 | Mauritian | Urine | control |
| CM24 | CM24 | Mauritian | Urine | control |
| CM25 | CM25 | Mauritian | Urine | control |
| CM26 | CM26 | Mauritian | Urine | control |
| CM27 | CM27 | Mauritian | Urine | control |
| CM28 | CM28.1 | Mauritian | Urine | control |
| CM29 | CM28.2 | Mauritian | Urine | control |
| CM30 | CM30 | Mauritian | Urine | control |
| CM31 | CM31 | Mauritian | Urine | control |
| CM32 | CM32 | Mauritian | Urine | control |
| CM33 | CM33 | Mauritian | Urine | control |
| CM34 | CM34 | Mauritian | Urine | control |
| CM35 | CM35 | Mauritian | Urine | control |
| CM36 | CM36 | Mauritian | Urine | control |
| CM37 | CM37 | Mauritian | Urine | control |
| CM38 | CM38 | Mauritian | Urine | control |
| CM39 | CM39 | Mauritian | Urine | control |
| CM40 | CM40 | Mauritian | Urine | control |
| CM41 | CM41 | Mauritian | Urine | control |
| CM42 | CM42 | Mauritian | Urine | control |
| CM43 | CM43 | Mauritian | Urine | control |
| CM44 | CM44 | Mauritian | Urine | control |
| CM45 | CM45 | Mauritian | Urine | control |
| CM46 | CM46 | Mauritian | Urine | control |
| CM47 | CM47 | Mauritian | Urine | control |
| CM48 | CM48 | Mauritian | Urine | control |
| CM49 | CM49 | Mauritian | Urine | HSCT-recipient |
| CM50 | CM50 | Mauritian | Urine | HSCT-recipient |
| CM51 | CM51 | Mauritian | Urine | HSCT-recipient |
| CM52 | CM52 | Mauritian | Urine | HSCT-recipient |
| CM53 | CM53 | Mauritian | Urine | HSCT-recipient |
| CM54 | CM54.1 | Mauritian | Urine | HSCT-recipient |
| CM55 | CM54.2 | Mauritian | Urine | HSCT-recipient |
| CM56 | CM56 | Mauritian | Urine | HSCT-recipient |
| CM57 | CM57.1 | Mauritian | Urine | HSCT-recipient |
| CM58 | CM57.2 | Mauritian | Urine | HSCT-recipient |
| CM59 | CM59.1 | Mauritian | Urine | HSCT-recipient |
| CM60 | CM59.2 | Mauritian | Urine | HSCT-recipient |
| CM61 | CM61.1 | Mauritian | Urine | HSCT-recipient |
| CM62 | CM61.2 | Mauritian | Urine | HSCT-recipient |
| CM63 | CM63 | Mauritian | Urine | HSCT-recipient |
| CM64 | CM64.1 | Mauritian | Urine | HSCT-recipient |
| CM65 | CM64.2 | Mauritian | Urine | HSCT-recipient |
| CM66 | CM64.3 | Mauritian | Urine | HSCT-recipient |
| CM67 | CM67.1 | Mauritian | Urine | HSCT-recipient |
| CM68 | CM67.2 | Mauritian | Urine | HSCT-recipient |
| CM69 | CM69.1 | Mauritian | Urine | HSCT-recipient |
| CM70 | CM69.2 | Mauritian | Urine | HSCT-recipient |
| CM71 | CM69.3 | Mauritian | Urine | HSCT-recipient |
| CM72 | CM72 | Philippine | Urine | control |
| CM73 | CM73 | Philippine | Urine | control |
| CM74 | CM74 | Philippine | Urine | control |
| CM75 | CM75 | Philippine | Urine | control |
| CM76 | CM76 | Philippine | Urine | control |
| CM77 | CM77 | Philippine | Urine | control |
| CM78 | CM78 | Philippine | Urine | control |
| CM79 | CM79 | Mauritian | Urine | HSCT-recipient |
| CM80 | CM80 | Mauritian | Urine | HSCT-recipient |
| CM81 | CM81.1 | Mauritian | Plasma | HSCT-recipient |
| CM81 | CM81.2 | Mauritian | Spleen | HSCT-recipient |

MafaPyV2 is one of the closest relatives to BKPyV known to date, with an 83% nucleotide identity. The absolute closest relative to MafaPyV2 is *Cercopithecus erythrotis* polyomavirus, with an 84.6% nucleotide homology (Fig. 2A). Genomic features of MafaPyV2 are characteristic of other polyomaviruses, including a 5,164 bp circular genome with early and late coding genes in opposite sense and a bidirectional promoter (noncoding control region) (Fig. 2B). Based on phylogenetic analysis of the nucleotide and protein sequence of VP1 from samples with complete coverage of VP1 (n=22), MafaPyV2 can be categorized into three discrete genotypes (type I-III) (Fig. 3A). Samples with detectable MafaPyV2 included both HSCT recipient and healthy control MCMs and there was no observable difference in genotype distribution between these two groups. MafaPyV2 was identified in 55% (32/58) of all cynomolgus macaques, with 75% of HSCT recipients and 48% of healthy animals shedding the virus (Fig. 4A-D and Table S1). Additionally, in HSCT recipients, MafaPyV2 was shed at significantly higher relative loads, with a 686-fold increase, compared to healthy controls. (Fig. 4E).

**Fig. 2.**
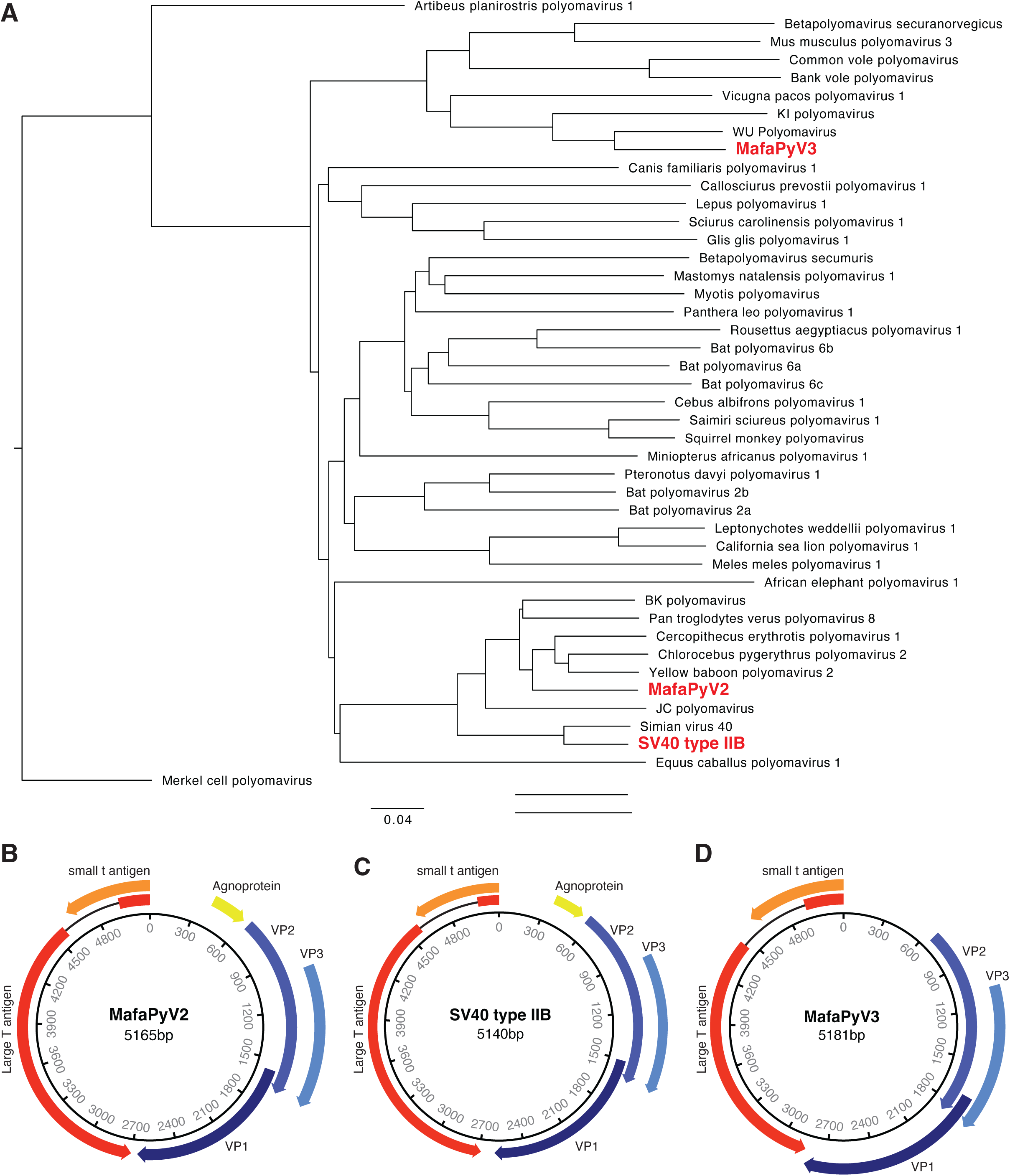
(A) Phylogenetic tree of Betapolyomavirus complete genome reference sequences and their relationship to MafaPyV2, MafaPyV3, and SV40 type IIB (bold in red). Merkel cell polyomavirus (an alphapolyomavirus) is serving as an outgroup. (B-D) Genome maps of (B) MafaPyV2, (C) MafaPyV3, and (D) SV40 type IIB.

**Fig 3.**
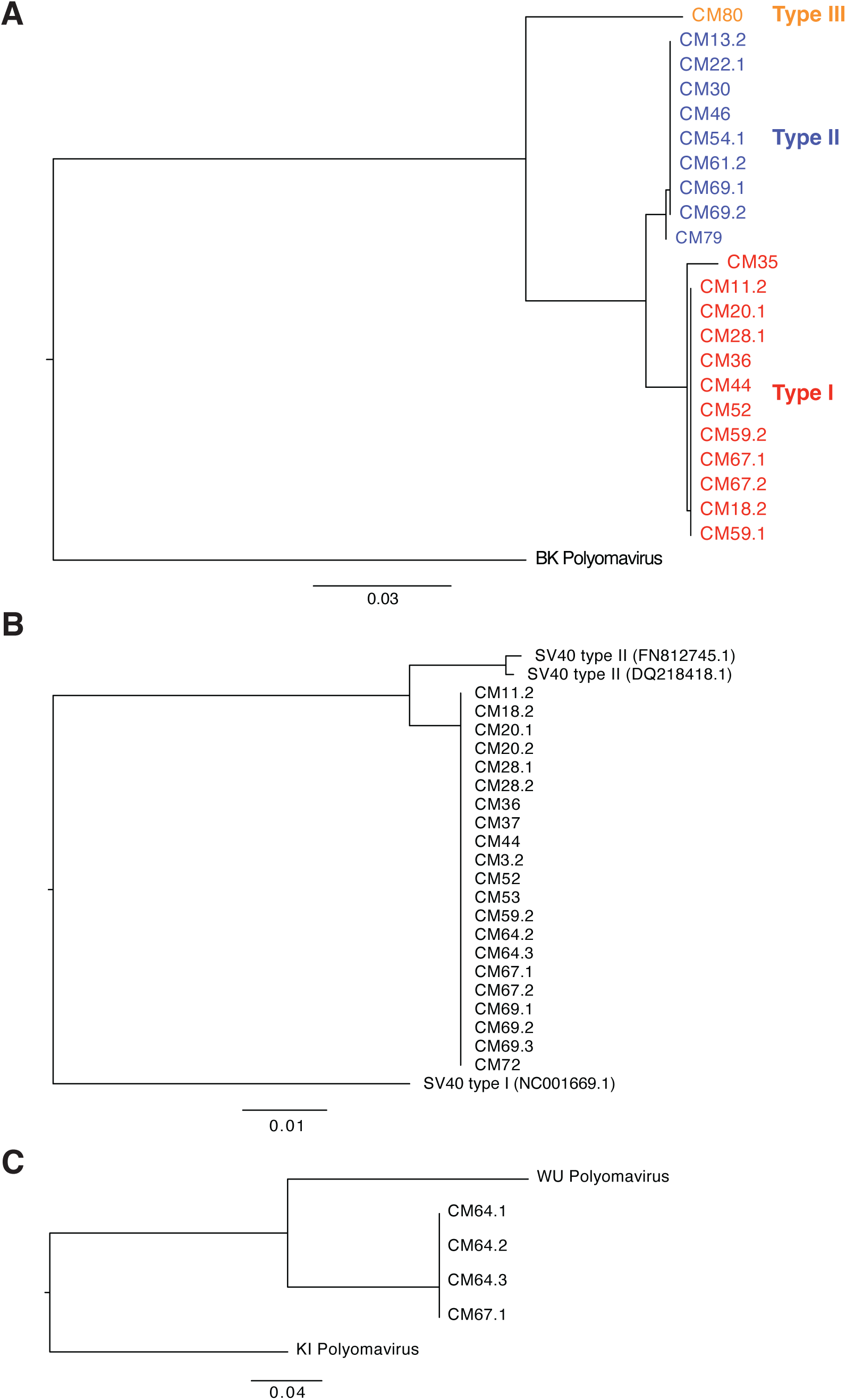
Phylogenetic tree of major capsid protein, VP1, nucleotide sequences from samples with complete coverage. (A) MafaPyV2. (B) SV40 type IIB. (C) MafaPyV3.

**Fig 4.**
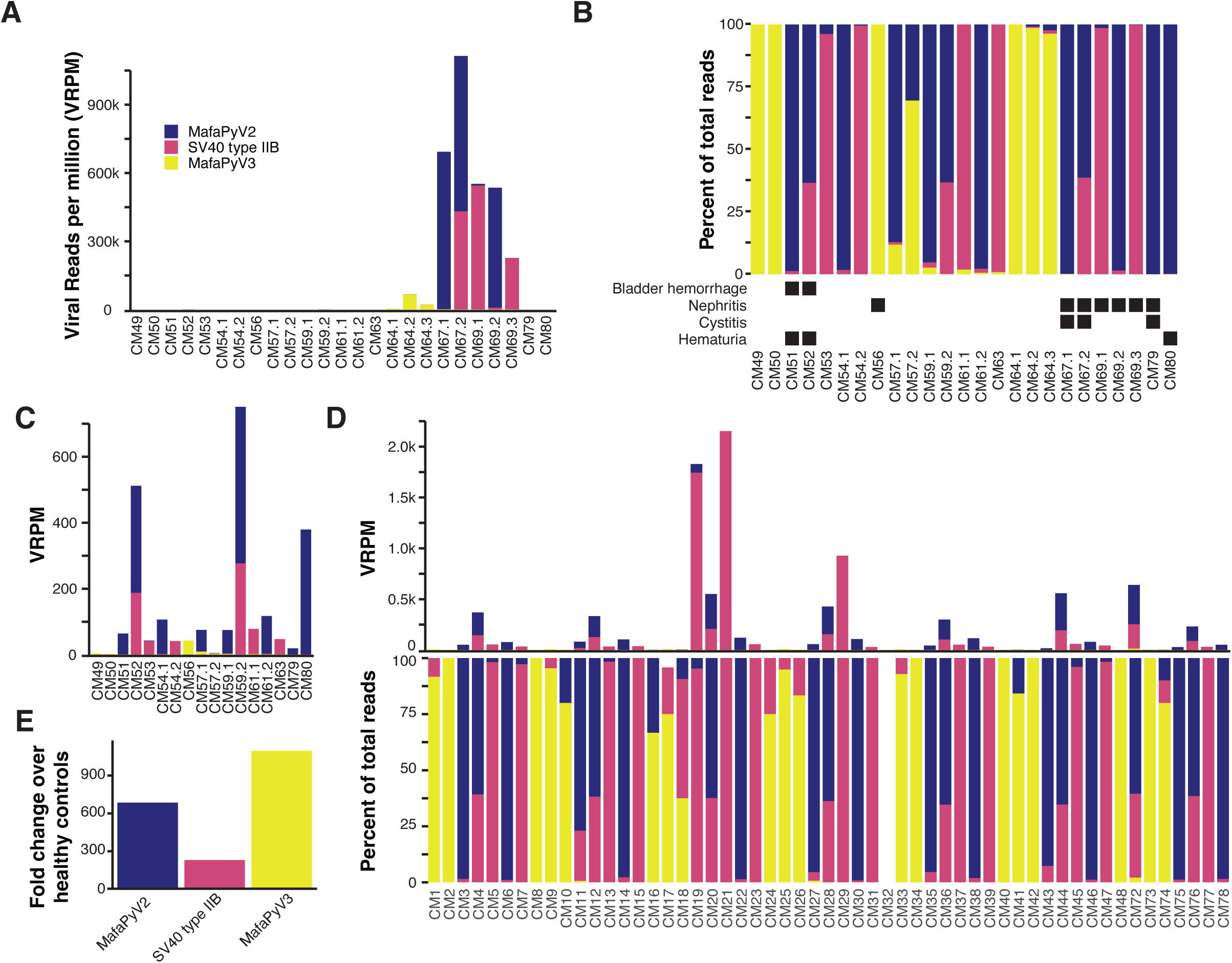
Polyomavirus species abundance in urine of HSCT-recipient cynomolgus macaques. (A) Abundance of polyomavirus species shed in urine of HSCT-recipients represented by viral reads per million (VRPM). (B) Percent of polyomavirus species reads of total polyomavirus reads for all HSCT-recipients with urinary clinical features indicated by black boxes. (C) Abundance of polyomavirus species shed in urine of HSCT-recipients represented by VRPM, excluding animals CM64, CM67, and CM69. (D) Polyomavirus shedding in healthy control cynomolgus macaques in VRPM and as percent of reads. (E) Raw fold difference in polyomavirus species reads between HSCT-recipient and healthy control cynomolgus macaques.

SV40 type IIB, a newly identified strain of SV40, is most closely related to SV40 type II, with 96.9% nucleotide identity (<u>DQ218418.1</u> and <u>FN812745.1</u>) (Fig. 3B). SV40 type II was first identified in 2011 in Simian/Human Immunodeficiency Virus-infected and healthy Chinese and Indian origin rhesus macaques^10^ and has an 88% nucleotide homology to SV40 type I, the classical type of SV40. Samples with complete coverage of SV40 type IIB VP1, including 14 Mauritian and one Cambodian origin CM, did not reveal any genomic variation between their consensus sequences (Fig. 3B). The homogenous genomic divergence from SV40 type II suggests this virus is a discrete lineage of SV40 type II, leading to its designation as type IIB. SV40 type IIB was found at similar prevalences between HSCT recipients (56%, 9/16) and healthy animals (59%, 25/42) (Fig. 4A-D and Table S1). However, HSCT recipients shed SV40 type IIB at relative loads more than 200-fold higher than healthy controls (Fig. 4E).

MafaPyV3 is most closely related to human WU polyomavirus, with an 82.2% nucleotide identity (Fig. 2A). Similar to its closest relatives, WU and KI polyomavirus, MafaPyV3 lacks an agnoprotein open reading frame (ORF) (Fig 2D). Although, a possible divergent sequence or spliced (ORF), as seen for HPyV7,^11^ may make identification difficult and the presence of an agnoprotein cannot be completely ruled out from this genomic data. Four complete MafaPyV3 genomes were assembled from two HSCT recipient MCMs with no genetic variation within VP1 (Fig. 3C). Overall, MafaPyV3 was shed at much lower relative loads compared to the other polyomaviruses identified. MafaPyV3 was identified in 34% (20/58) of cynomolgus macaques, with 56% (9/16) of HSCT recipients and 26% (11/42) of healthy animals shedding the virus (Fig. 4A-D and Table S1). In HSCT recipients, MafaPyV3 was shed at significantly higher relative loads, exceeding a 1,000-fold increase, compared with healthy controls (Fig. 4E). In the urine of the two Indonesian-origin cynomolgus macaques, only MafaPyV3 was identified, and in the two Cambodian-origin cynomolgus macaques both MafaPyV2 and SV40 type IIB were found. Among urine samples from Philippine-origin cynomolgus macaques, all three polyomaviruses were present, with MafaPyV2 most frequently detected and MafaPyV3 least frequently detected. All three polyomaviruses were detected in the urine of HSCT recipient and healthy MCMs (Table S1).

### Urologic disease and associated polyomaviruses in HSCT recipient Mauritius cynomolgus macaques

Seven HSCT recipient MCMs with urine samples presented with urinary clinical signs and/or urinary post-mortem diagnoses after transplantation. Clinical signs included macroscopic hematuria or microscopic hematuria on urinalysis and post-mortem findings included nephritis (+/- large T antigen immunoreactivity) and/or cystitis (+/-hemorrhagic) (Table 2). Additionally, these animals frequently exhibited anorexia and weight loss and two animals with urine samples had confirmed viremia or viruria antemortem. Seven HSCT recipients developed acute cutaneous and/or multiorgan GVHD (Table S1).

**Table 2.**

| Animal ID | Urologic clinical signs/disease | Predominant polyomaviruses |
| --- | --- | --- |
| CM51 | Bladder hemorrhage | MafaPyV2 |
| CM52 | Microscopic hematuria, bladder hemorrhage | MafaPyV2 and SV40 type IIB |
| CM56 | Microscopic hematuria, chronic interstitial nephritis and cystitis (SV40 LT IHC negative) | MafaPyV3 |
| CM67 | Cystitis and PyV-associated nephritis | MafaPyV2 and SV40 type IIB |
| CM69 | Mild renal tubular injury and chronic interstitial nephritis | MafaPyV2 and SV40 type IIB |
| CM79 | Mild nephritis and cystitis | MafaPyV2 |
| CM80 | Antemortem hematuria (alive) | MafaPyV2 |
| CM81<br>(spleen) | Hemorrhagic cystitis and PyV-associated nephritis | SV40 type IIB |

All seven animals with urologic disease were shedding one or multiple polyomaviruses (Fig. 4B). Two animals (CM67 and CM69) had exceptionally high relative viral loads of MafaPyV2 and SV40 type IIB and one of those animals (CM67) had confirmed polyomavirus-associated nephritis and cystitis (Fig. 4A-B, and Table 2). This animal was shedding all three polyomaviruses in the urine with predominance of MafaPyV2 and minimal shedding of MafaPyV3. The other animal (CM69) had mild renal tubular injury and nephritis and was also shedding all three polyomaviruses but with a predominance of SV40 type IIB and minimal shedding of MafaPyV3 (Fig. 4A-B, and Table 2). The five other animals with urologic signs/disease had mild bladder hemorrhage, mild nephritis and cystitis or mild renal tubular injury and disease did not correspond to high levels of polyomavirus shedding (Fig. 4A-C). The healthy control animals were shedding a lower relative abundance of polyomavirus overall, with a predominance of SV40 type IIB (Fig. 4D).

We also evaluated one plasma and spleen sample from an additional HSCT recipient MCM, without an available urine sample, that was diagnosed with severe hemorrhagic cystitis and polyomavirus-associated nephritis postmortem. Short read and long read sequencing performed on the plasma DNA did not identify any polyomaviruses, likely due to low DNA concentration. However, short read and long read sequencing on extracted spleen DNA revealed the sole presence of SV40 type IIB, although no large T antigen immunoreactivity was detected in the spleen tissue (Table S1).

### Inter and intra-host variants of polyomaviruses in cynomolgus macaques

To assess the genetic diversity of the polyomaviruses in the urine within and between animals, we performed variant calling analysis using LoFreq and annotation of coding regions using snpEff. Variants of MafaPyV2 occurred within the noncoding control region (NCCR) and all coding regions. Most inter-host variants were synonymous substitutions with 65 unique synonymous variants, while missense variants were less common with 20 unique missense variants (Table S4). Unique missense variants occurred most commonly within the agnoprotein (19.0 variants/kb), VP2 (4.7 variants/kb), and VP1 (3.7 variants/kb) when normalized by gene length. Within VP1 of MafaPyV2, three inter-host missense variants separated genotype II from genotype I (Fig. 5A and B). One surface variant represented by a G>T mutation in residue 64 resulting in a glutamic acid to aspartic acid substitution was three times more frequent in the urine of HSCT recipients (n_HSCT_=3, n_Control_=1). While the other two missense variants were present in the urine of both HSCT recipients and healthy controls, at similar frequencies, and included a A>C mutation in residue 188 resulting in a neutral glutamic acid to aspartic acid substitution on the VP1 surface and a G>T mutation in residue 156 resulting in an alanine to serine substitution within the inner beta sheet. One MafaPyV2 strain within genotype I clade (CM67.2), from an animal with polyomavirus-associated nephritis, exhibited a low frequency (0.008 AF) G>A mutation in residue 188 resulting in a glutamic acid to lysine amino acid substitution and a charge reversal from negative to positive. The intra-host missense variants of VP1 in MafaPyV2 overlapped with these 4 described inter-host variants (Table S7).

**Fig 5.**
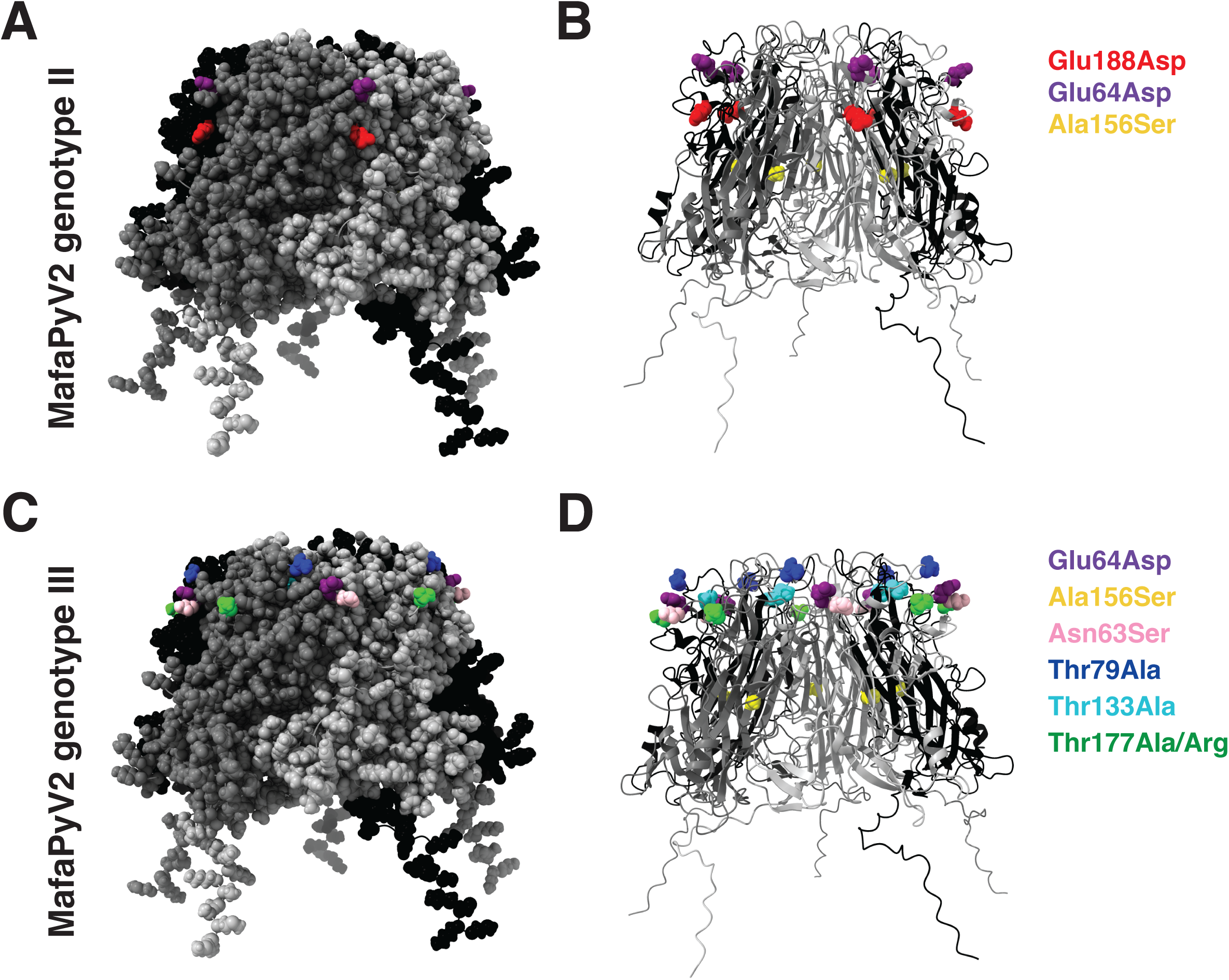
MafaPyV2 VP1 missense variants. (A) MafaPyV2 genotype II missense variants relative to genotype I highlighted on the VP1 pentamer with space filling atom and (B) ribbon structures, Glu188Asp in red, Glu64Asp in purple and Ala156Ser in yellow. Variants were identified using short read sequencing data. (C) MafaPyV2 genotype III (CM80) missense variants relative to genotype I highlighted on the VP1 pentamer with space filling atom and (D) ribbon structures. Two missense variants resulted in the same amino acid changes present in the genotype II, Glu64Asp in purple and Ala156Ser in yellow, and five additional unique missense variants resulting in four amino acid changes, Asn63Ser in pink, Thr79Ala in blue, Thr133Ala in cyan, and Thr177Ala/Arg in green. Variants were identified using long read sequencing data from CM80.

Low sequencing depth and coverage from Illumina shotgun sequencing of CM79 and 80 restricted the ability to detect variants in MafaPyV2 through LoFreq. Therefore, alignment, variant calling and annotation were performed on the long read sequences of MafaPyV2 using minimap2, bcftools and snpEff (Table S5-6). MafaPyV2 from CM80 represents the most divergent genotype (type III) and compared to genotype I is characterized by six missense variants in VP1, including two missense variants that result in the same amino acid change present in genotype II, Glu64Asp and Ala156Ser, and five additional missense variants resulting in four amino acid changes, Asn63Ser, Thr79Ala, Thr133Ala, Thr177Ala, and Thr177Arg (Fig 5C-D). CM80 was isolated from an animal 5 months after intermittent hematuria began. This animal did not succumb to disease and remains alive. The only missense variants detected in VP1 of MafaPyV2 from CM79 were the three missense variants that distinguish genotype II from genotype I in VP1.

For Large T antigen, the inter and intra-host missense variants of MafaPyV2 overlapped and included 7 variants within 16 samples from 13 animals. Six variants occurred in both HSCT recipients and healthy controls, and one variant only occurred in one healthy control (Table S4 and S7). An additional 13 missense variants within Large T antigen of MafaPyV2 were identified from CM79 and CM80 (Table S5-6).

There were only three inter-host missense variants observed in SV40 type IIB, all occurring within Large T antigen and identified in a healthy control and an HSCT recipient (Table S8-9). There were no variants identified in SV40 type IIB identified in the spleen of CM81. All variants within MafaPyV3 occurred in HSCT recipients and included one missense variant within VP1 and VP2 (Table S10-11). Additionally, occasional indels occurred throughout the three polyomavirus genomes that resulted in frameshift or nonsense mutations.

### Longitudinal intra-host polyomavirus genetic variability

To determine if acute virus evolution can be observed, longitudinal urine samples were evaluated from seven HSCT recipient MCMs and 12 healthy cynomolgus macaques, with 2-3 timepoints from each animal. Multiple polyomaviruses were commonly shed together in a single urine sample and shedding of polyomaviruses was occasionally intermittent between longitudinal samples. Out of 19 animals with multiple urine collection timepoints, 10, 5, and 4 animals were intermittently shedding MafaPyV2, SV40 type IIB, and MafaPyV3 respectively (Table S1). Only three HSCT recipients had detection and adequate coverage of the polyomaviruses at multiple timepoints for inter-host variant analysis. Sample 67.2, coinciding with the detection of polyomavirus viruria and viremia, revealed a unique missense variant (0.008 AF), Glu188Lys, of MafaPyV2 on a surface-exposed helix of VP1 that is not present at an earlier timepoint (CM67.1). Another animal, CM69 revealed substantial variability between CM69.1-2 and CM69.3 (necropsy sample) for MafaPyV2, placing them in separate clades on the complete genome phylogenetic tree (Figure S3). However, no missense variants of VP1 were identified between them due to low coverage of VP1 in CM69.3. This possibly suggests either a new infection with another virus strain or expansion of an existing genotype I co-infection at this later timepoint instead of intra-host evolution. Samples from CM64 revealed no detection of MafaPyV2 at an early timepoint (CM64.1; 29 days post-HSCT) but intra-host missense variants differed between CM64.2 and .3, which were two aliquots from a single collected sample (52 days post-HSCT). This included one additional missense variant in VP1 and 3 additional missense variants in Large T antigen of CM64.3 and one unique missense variant in VP2 for both CM64.2 and .3. Additionally, there was one missense variant within Large T antigen in CM64.2 that was not present in CM64.3 (Table S4).

### Cynomolgus macaque polyomaviruses show evidence of long term APOBEC3 selection pressure

Many human DNA viruses, including polyomaviruses, exhibit genomic signatures consistent with long-term selection pressure imposed by host APOBEC3 cytosine deaminases^12^. These signatures are thought to arise through prolonged virus-host coevolution, resulting in characteristic depletion of APOBEC3 target dinucleotides. We sought to determine whether these polyomaviruses identified here display analogous patterns of APOBEC3-associated selection. To investigate whether APOBEC3 has had an impact on these cynomolgus macaque virus genome compositions over a long evolutionary time frame, we calculated the enrichment of TC/TT and GA/AA dinucleotides in their genomes by observed versus expected by Markov modeling. TC and GA dinucleotides represent the 3’ cytosines on the positive and negative strand that can be targeted by APOBEC3 enzymes and TT and AA are the resulting products of deamination and DNA replication. All three genomes exhibited a depletion of TC:GA dinucleotides and an overrepresentation of TT:AA dinucleotides (Fig. 6A-C).

**Fig 6.**
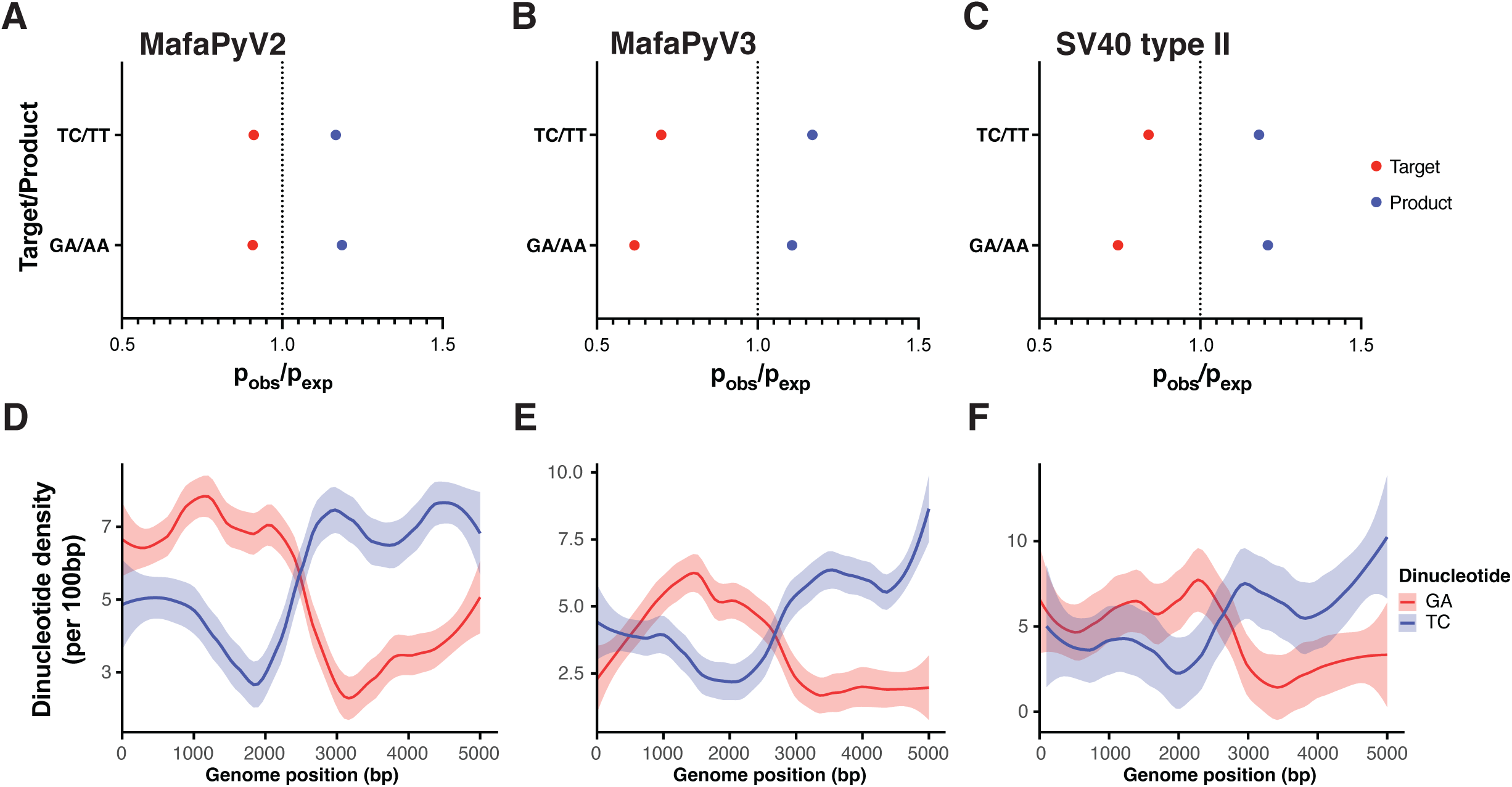
Evidence of long-term APOBEC3 selection pressure on the cynomolgus macaque polyomavirus genomes. (A) Representation of potential APOBEC3 target and product dinucleotides in MafaPyV2 (B) MafaPyV3 (C) and SV40 type IIB. (D) Line graph of TC/GA dinucleotide density using 100-bp nonoverlapping windows and smoothed fitted lines with 95% confidence intervals of these densities across the MafaPyV2, (E) MafaPyV3, and (F) SV40 type IIB genomes, starting at the late coding region.

Additionally, we observed an inverse relationship between TC and GA dinucleotide density on these cynomolgus macaque polyomavirus genomes that corresponds to replication and transcription direction (Fig. 6D-F).

To assess potential ongoing mutational processes on these viruses, we determined the trinucleotide contexts of intra-sample variants. In MafaPyV2, most mutations occurred at CpG sites when normalized by trinucleotide abundance (Fig. S2A). Within the MafaPyV3 genome, only C>T and C>G mutations at TCA, TCT, and CCG trinucleotides were observed, potentially representing an APOBEC3 mutation signature (Fig. S2B). However, the significance of this signature is unclear with such low variant frequency. For SV40 type IIB, there was no APOBEC3-mediated mutation signature (Fig. S2C).

## Discussion

In this study we broadly evaluated the DNA virome and comprehensively characterized the genetics of the polyomaviruses present in the urine of immunocompromised and healthy cynomolgus macaques using complementary short- and long-read sequencing approaches. We identified three betapolyomaviruses with apparent specific tropism to cynomolgus macaques: *Macaca fascicularis* polyomavirus 2 (MafaPyV2), *Macaca fascicularis* polyomavirus 3 (MafaPyV3), and a newly described strain of simian virus 40, SV40 type IIB. Importantly, both MafaPyV2 and SV40 type IIB were previously identified as “cynomolgus polyomavirus” (CPV), and our complete genomic analyses resolve this ambiguity by demonstrating that they represent distinct viruses.

In our cohort, although all three polyomaviruses were detected in both immunocompromised and immunocompetent macaques, HSCT recipients exhibited dramatically higher relative viral loads, exceeding several hundred- to thousand-fold increases, compared to control animals. This pattern closely parallels observations in human HSCT recipients, where immunosuppression permits reactivation and uncontrolled replication of latent polyomaviruses, such as BKPyV. All HSCT recipients that developed clinical or histopathologic evidence of urologic disease were shedding one or more polyomaviruses, and multiple polyomaviruses were frequently detected within a single host. The two animals with the most severe disease phenotypes exhibited exceptionally high relative loads of either MafaPyV2 and/or SV40 type IIB, supporting a model in which viral burden contributes to disease severity, similar to the strong association between high BKPyV viruria and viremia and hemorrhagic cystitis or nephropathy in humans^13,14^. In contrast, animals with milder urologic lesions generally exhibited lower viral loads, suggesting that polyomavirus detection alone is insufficient to predict disease outcome. Because this study relied on retrospective antemortem urine sampling and postmortem diagnoses, these findings do not establish a causal relationship between individual polyomaviruses and specific disease phenotypes.

The polyomavirus major capsid protein, VP1, is the principal determinant of cellular surface receptor binding and the primary target of the host antibody response^4^. In this study, we identified several missense variants occurring at surface-exposed or structurally important regions. In human BKPyV, analogous VP1 polymorphisms influence receptor usage, cell tropism, and susceptibility to neutralizing antibodies, raising the possibility that MafaPyV2 VP1 variants may similarly modulate viral fitness or immune evasion^15^. Notably, certain VP1 variants of MafaPyV2 were more frequently observed in HSCT recipients than in healthy controls, suggesting that immune suppression may permit the expansion or persistence of viral subpopulations with altered functional properties as in humans. However, given the limited number of animals and variable sequencing depth across samples, these interpretations remain speculative. Functional studies examining receptor binding, antibody neutralization, and cell-type specificity would need to be performed to determine whether these variants confer biologically meaningful differences in viral behavior. In contrast, SV40 type IIB and MafaPyV3 exhibited limited genetic diversity. The low diversity of SV40 type IIB may reflect either limited mutagenic processes active on the virus or stronger host constraints causing any variant to have a substantial decrease in fitness.

All three cynomolgus macaque polyomaviruses exhibited signatures of long-term APOBEC3-mediated selection pressure, including depletion of TC/GA dinucleotides and enrichment of TT/AA dinucleotides across their genomes. This pattern mirrors findings in human polyomaviruses and supports a conserved role for APOBEC3 enzymes in shaping polyomavirus genome composition over evolutionary timescales^12,16^. In contrast to reports in human disease, evidence for APOBEC3-mediated mutagenesis at the level of intra-host viral variation was limited and inconsistent. This suggests that while APOBEC3 activity has exerted a measurable influence on viral evolution over long periods, it may contribute only sporadically to mutational processes during active infection. This may also be explained by macaque APOBEC3B being cytoplasmic and having no or limited access to the viral chromatin during infection^17^. Therefore, the apparent APOBEC3-target depleted genome composition may represent an ancestral bottleneck rather than an active process on these viruses, but further studies are required to determine this.

Polyomaviruses, especially those belonging to the genus Beta, are largely considered to slowly evolve alongside their host with limited interspecies transmission^18^. The Rhesus-Cynomolgus macaque divergence is estimated to be about 0.9 Mya^19^ with Rhesus genetic introgression into the CM genome occurring more frequently in Indochinese CM, suggesting more recent interbreeding/interactions. Based on the branch length between BKPyV and its next closest relative *Pan troglodytes verus* polyomavirus and assuming these viruses diverged around the same time as the divergence time between *P. troglodytes verus* and humans (5-6 Mya)^20^, SV40 type IIB divergence from SV40 type I likely occurred 2.7-4.2 Mya and diverged from SV40 type II between 0.56 to 0.81 Mya. Collectively, these data indicate that SV40 type IIB arose after the speciation of CM and, in light of its absence in rhesus macaques, is likely a CM-specific virus.

Cynomolgus macaques have a native range spanning from southern China, eastern India, through Southeast Asia and the Indonesian archipelago. A founding population of an estimated 10-20 cynomolgus macaques were deposited on the island of Mauritius around the 1600s and has expanded to over 30,000 animals living on the island^19,21^. This significant bottleneck has been capitalized on for immunology research using the limited genetic diversity of these animals^7^, but it also represents an opportunity for geographically isolated viral divergence. We specifically included macaques from various geographic origins to address whether any viruses, strains, or lineages were specific to any geographic location. While this was not observed potentially reflecting the slow evolutionary dynamics of polyomaviruses, it may also be due to the captive rearing and co-housing of animals or low shedding in control animals. Whether the diversity and infection frequency observed in this cohort of animals applies to wild animals would need further study.

Metagenomic analyses of urine in human HSCT recipients remain limited, with clinical monitoring largely restricted to targeted approaches such as PCR assays for BKPyV, JC polyomavirus, and adenovirus^22^. However, shotgun metagenomic analyses of urine from renal transplant patients have revealed diverse viromes that include polyomaviruses, papillomaviruses, anelloviruses, and herpesviruses^22,23^. In immunocompetent humans, however, the urinary virome is typically dominated by bacteriophages, with comparatively fewer reads corresponding to eukaryotic DNA viruses such as polyomaviruses, papillomaviruses, anelloviruses, and herpesviruses^24,25^. These findings closely mirror the virome composition identified in this cynomolgus macaque cohort and these parallels underscore the translational relevance of the MCM HSCT model for studying opportunistic viral infections in immunocompromised hosts.

The management of infectious disease complications, specifically BKPyV-associated disease, after allogeneic HSCT continues to be challenging due to the lack of specific antivirals and preventative measures. Approximately 10-29% of allogeneic HSCT patients develop BKPyV-associated hemorrhagic cystitis after engraftment and disease is associated with high BKPyV load in the urine^3,26,27^. Furthermore, development of high grade hemorrhagic cystitis is associated with a decreased overall survival and an increased risk of transplant-related mortality^27^. The identification of analogous viral reactivation and urologic disease in MCMs provides a valuable experimental system that cannot be recapitulated in mice. Murine cells are non-permissive for productive BKPyV infection and while mouse polyomavirus has been widely used in studies, it is an alphapolyomavirus with a distinct oncogenic driver (middle T antigen) and dissimilar pathogenesis^28,29^. Mice also only encode a single APOBEC gene and lack the diversity of APOBEC3 enzymes seen in primates^30^. Overall, this study characterized the polyomaviruses and broader virome in urine from cynomolgus macaques and establishes the Mauritian cynomolgus macaque HSCT model as a translational platform to study polyomavirus reactivation, host immune responses, viral evolution, and intervention strategies relevant to human transplantation.

## Methods

### Animals

Mauritian-origin cynomolgus macaques (*Macaca fascicularis)* were housed at the ONPRC and utilized for studies under the approval of the Oregon Health and Science University (OHSU) West Campus Institutional Animal Care and Use Committee (IACUC) under protocols #IP00000838. All macaques in this study were managed according to the ONPRC animal care program, which is fully accredited by AAALAC International and is based on the laws, regulations, and guidelines set forth by the United States Department of Agriculture (e.g., the Animal Welfare Act and Animal Welfare Regulations, the Guide for the Care and Use of Laboratory Animals, 8^th^ edition (Institute for Laboratory Animal Research), and the Public Health Service Policy on Humane Care and Use of Laboratory Animals. The nutritional plan utilized by the ONPRC is based on National Research Council recommendations and supplemented with a variety of fruits, vegetables, and other edible objects as part of the environmental enrichment program established by the Behavioral Services Unit. Animals were socially housed, when possible. All efforts were made to minimize suffering through the use of minimally invasive procedures, anesthetics, analgesics, and environmental enrichment. Transplanted recipient macaques included both SIV-naïve and SIV-infected macaques on antiretroviral therapy with fully suppressed SIV viremia.

### Sample and tissue procurement

80 urine samples were collected from 58 cynomolgus macaques (50 Mauritian, 6 Philippine, 1 Cambodian, and 1 Indonesian-origin macaque). 16 Mauritian cynomolgus macaques (MCMs) were immunosuppressed and samples were collected between 2 to 186 days after HSCT. The remaining 42 animals were healthy controls. Additionally, plasma and spleen homogenate samples were collected from one immunosuppressed MCM at day 45 post-HSCT. Urine was collected via pan catch or cystocentesis and clarified by centrifugation at 830rcf for 4 minutes. Plasma was isolated by centrifugation of whole blood (collected in EDTA tubes) at 1860rcf for 10 minutes. Plasma was removed and clarified at 830rcf for 4 minutes. Spleen was collected at necropsy in R10 media (RPMI1640, 10% fetal bovine serum, antibiotic/antimycotic (HyClone), 2mM L-glutamine, 1mM sodium pyruvate), diced and mashed through 70-micron cell strainers to collect single-cell suspension, and ACK-treated to lyse red blood cells.

### Urine and plasma nucleic acid isolation

100 µL of 5x Digestion Buffer (125 mM Tris pH 8.0, 125 mM EDTA, 50 mM DTT, 5% SDS, 2.5% proteinase K (Qiagen #19131)) was added to 400 µL of each sample. Samples were digested for 1 hour at 65°C. 1633 µL of buffer PM and 33 µL of 3M sodium acetate were added to each sample, which were then centrifuged in DNA spin columns at 16,000g for 1 minute. The spin columns were washed twice with 750 µl of PE buffer and then spun for 5 minutes at room temperature to achieve complete dryness. Samples were eluted in 30 µl of 0.1x TE (2 mM Tris pH 8, 0.2 mM EDTA) for five minutes at 65°C. Following centrifugation, the elution was transferred back to the spin column, and the incubation and centrifugation steps were repeated (this final elution will henceforth be referred to as the “unamplified sample”). DNA was quantified by Qubit and samples were stored at -20°C.

### Spleen cell homogenate nucleic acid isolation

Low-molecular-weight DNA isolation was performed with a modified HIRT extraction protocol. Up to 100 ul of PBS and 100 ul of 2x HIRT lysis buffer were added to a volume of 500,000 cells and incubated at room temperature for 20 minutes. 80 ul of 5M NaCl was added and incubated at 4°C overnight. 8ul of proteinase K and 4 ul RNAse A as added and incubated at 37°C for 1 hour. 1500 ul of Buffer PB was added and transferred to a DNA purification column, following the QIAquick PCR Cleanup protocol (Qiagen) and eluting with 30 ul H20 (this final elution will also be referred to as the “unamplified sample”). DNA was quantified by spectrophotometry and samples were stored at -20°C.

### Rolling circle amplification (RCA)

A 5 or 2.5 µL aliquot of the unamplified sample combined with 5 or 2.5 ul sample buffer, respectively, was heated at 95°C for 3 minutes before being returned to room temperature. 10 or 5 µl of TempliPhi RCA enzyme mixture was added to each heated RCA sample, mixed and then spun down briefly to reconsolidate. Samples were incubated for 6-24 hours at 30°C, then heat inactivated at 65° for 10 minutes. These samples are referred to as “RCA samples.” DNA was quantified by Nanodrop and Qubit.

### PCR amplification of MafaPyV2 and SV40 type IIB and nanopore long-read sequencing

The urine RCA samples were used as template DNA for PCR amplification of MafaPyV2.

Two sets of primers to cover the complete polyomavirus genome were used; Primer set #1 and 2 (Table S3). The spleen and plasma RCA samples from macaque CM80 were used as template DNA for PCR amplification of SV40 type IIB. Two sets of primers to cover the complete genome were used; Primer set #3 and 4 (Table S3).

### Short-read sequencing

Illumina Nextera tagmentation sequencing on the Illumina MiSeq (CM79 and 80) and NextSeq 2000 was performed by the CCR Genomics core on all RCA amplified urine and plasma samples.

### Bioinformatic Analysis

#### Sequence alignment

Illumina sequencing reads were trimmed using fastp 0.24.0 with default settings. Trimmed reads were aligned with Bowtie2 (2-2.5.3) to the cynomolgus macaque host genome and removed.

#### De novo assembly and annotation

All reads not mapping to the host genome were de novo assembled using MEGAHIT (1.2.9) with default parameters^31^. Assembled contigs were annotated using BLASTn and DIAMOND against the NCBI database for closely related species, and Cenote-Taker2 with default parameters was used to identify more divergent species^32^. A sample was considered positive for a polyomavirus if it had at least 8 reads aligned to the viral reference genome with Bowtie2.

#### Point mutation variant calling

Genomic variants of cynomolgus macaque poylomaviruses were called using LoFreq (version 2) and bcftools (samtools 1.21). Variants with amino acid changes were identified with SnpEff (version 5.2). Alphafold (version 2.3.2) was used to predict the structure of the VP1 pentamer for MafaPyV2. Inter and intra-host variants were mapped to the predicted structure.

#### Viral motif enrichment and density and mutation signature analysis

Dinucleotide and trinucleotide motif enrichments for each aligned genome were calculated using Markov modeling as described by Ebrahimi, et al^33^. Dinucleotide density was calculated across the viral genomes using 100-bp nonoverlapping windows. Smoothed fitted lines and 95% confidence intervals of these densities were calculated. Intra-sample variants were characterized by their trinucleotide contexts and single nucleotide substitution type. All mutations detected per virus were normalized by their respective reference motif abundance and visualized using R.

### Phylogenetic Analysis

Phylogenetic trees were generated using MAFFT version 7 and FigTree version 1.4.4.

### Data visualization

All graphs were made in R statistical environment (4.5.1) using the package ggplot2 or using GraphPad Prism (10.6).

Coding was assisted using AI tools. All code and output were carefully reviewed for accuracy.

## Data availability

All raw sequencing will be available in SRA upon publication of the final version of the manuscript.

Annotated complete genomes will be available in GenBank upon publication of the final version of the manuscript.

Code used in the bioinformatic analysis can be found at https://github.com/VogelHelena/Cyno_Virus_Discovery

## Credit authorship contribution statement

**Helena Vogel:** Investigation, Data curation, Formal analysis, Visualization, Writing – original draft, Writing – review & editing

**Tristan Neal:** Investigation, Data curation, Writing – review & editing

**Helen Wu:** Resources, Writing – review & editing

**Paul Kievit:** Resources, Writing – review & editing

**Jonah Sacha:** Resources, Writing – review & editing

**Gabriel Starrett:** Conceptualization, Formal analysis, Visualization, Data curation, Writing – original draft, Writing – review & editing, Supervision.

## Acknowledgments

The laboratory of GJS is supported by the Center for Cancer Research, National Cancer Institute, National Institutes of Health Intramural Research Program project number ZIA BC 011894. The contributions of the NIH author are considered works of the United States Government. The findings and conclusions presented in this paper are those of the authors and do not necessarily reflect the views of the NIH or the U.S. Department of Health and Human Services. Helena Vogel is in the NIH Comparative Biomedical Scientist Training Program supported by the National Cancer Institute in partnership with Colorado State University. We would like to thank Chris Buck, Diana Pastrana, and the rest of the Laboratory of Cellular Oncology for their thoughtful discussions on the manuscript. The CCR Genomics core provided sequencing services. This work utilized the computational resources of the NIH HPC Biowulf cluster (https://hpc.nih.gov). The work at the Oregon National Primate Center was supported by National Institutes of Health grants R21 AI112433, R37AI189285, and R01 AI129703 awarded to JBS.

## Supplemental material

Fig. S1. Partial amino acid alignment of Large T antigen between previously isolated samples and MafaPyV2 and SV40 type IIB.

Fig. S2. Normalized intra-host mutational signatures of A) MafaPyV2, B) MafaPyV3, and C) SV40 type IIB.

Fig. S3. Phylogenetic tree of MafaPyV2 complete genomes with at least 90% coverage.

Table S1. Cynomolgus macaque sample collection with clinical and sequencing data.

Tabe S2. Viral group abundance in HSCT-recipient and healthy control cynomolgus macaques. Viral reads are normalized per sample to reads per million before combining reads.

Table S3. PCR primers for long-read sequencing.

Table S4. MafaPyV2 inter-host variants. (All.Mafapyv2.interhost.variants.shortread.xlsx)

Table S5. CM79 MafaPyV2 inter-host variants. (MafaPyV2CM79.bcftools.longread.xlsx)

Table S6. CM80 MafaPyV2 inter-host variants. (MafaPyV2CM80.bcftools.longread.xlsx)

Table S7. MafaPyV2 intra-host variants. (All.MafaPyV2_intrahost.ann.vars.xlsx)

Table S8. SV40 type IIB inter-host variants (All.SV40typeIIB.interhostvariants.shortread.xlsx)

Table S9. SV40 type IIB intra-host variants. (All.SV40typeIIB.intrahost.ann.vars.xlsx)

Table S10. MafaPyV3 inter-host variants. (All.Mafapyv3.interhost.variants.shortread.xlsx)

Table S11. MafaPyV3 intra-host variants. (All.MafaPyV3.intrahost.ann.vars.xlsx)

