## Supplementary material for "Characterization of the urinary DNA virome of hematopoietic stem cell transplant recipient and healthy cynomolgus macaques": Figure S1

### Fig. S1

| | | | | | | | | | | | | | | | | | | | | | | | | | | | | | | | | | | | | | | | | | | | | | |
|---|---|---|---|---|---|---|---|---|---|---|---|---|---|---|---|---|---|---|---|---|---|---|---|---|---|---|---|---|---|---|---|---|---|---|---|---|---|---|---|---|---|---|---|---|---|
| SV40 type I | . | . | . | . | . | . | . | . | . | . | . | E | - | - | - | . | N | . | . | . | . | . | . | . | . | . | . | . | D | . | . | . | . | . | . | . | . | . | . | . | . | . | . | . | . |
| SV40 type II | . | . | . | . | . | . | . | . | . | . | . | - | - | - | - | . | . | . | . | . | . | . | . | . | . | . | . | . | . | . | . | . | . | . | . | . | . | . | . | . | . | . | . | . | . |
| SV40 type IIB | . | . | . | . | . | . | . | . | . | . | . | - | - | - | - | . | . | . | . | . | . | . | . | . | . | . | . | . | . | . | . | . | . | . | . | . | . | . | . | . | . | . | . | . | . |
| SV40 type IIB (CM81) [33460 (Wu et al., 2019)] | . | . | . | . | . | . | . | . | . | . | . | - | - | - | - | . | . | . | . | . | . | . | . | . | . | . | . | . | . | . | . | . | . | . | . | . | . | . | . | . | . | . | . | . | . |
| "CPV" (Gorder et al., 1999) | . | . | . | . | . | . | . | . | . | . | . | - | - | - | - | . | . | . | . | . | . | . | . | . | . | . | R | . | . | . | . | . | . | . | . | . | . | . | . | . | . | . | R | E | K |
| BKPyV | E | . | . | . | S | . | . | S | S | . | . | E | . | . | . | . | . | . | . | . | H | . | D | . | F | A | . | . | . | . | . | . | . | . | . | . | . | . | . | . | . | . | . | . | . |
| MafaPyV2 (CM28) | E | . | . | . | A | . | . | . | . | . | . | D | . | . | . | . | E | . | . | . | H | . | D | . | F | A | . | . | . | . | . | . | . | . | . | . | . | . | . | . | . | . | . | . | K |
| MafaPyV2 (CM80) [33459 (Wu et al., 2019) | E | . | . | . | A | . | . | . | . | . | . | D | . | . | . | . | E | . | . | . | H | . | D | . | F | A | . | . | . | . | T | . | . | . | . | . | . | . | . | . | . | . | . | . | K |
| consensus | D | E | W | E | Q | W | W | N | A | F | N | | K | W | D | E | D | L | F | C | S | E | E | M | P | S | S | D | E | E | A | T | A | D | S | Q | H | S | T | P | P | K | K | K | R |
