## Supplementary figures and images for "Characterization of the urinary DNA virome of hematopoietic stem cell transplant recipient and healthy cynomolgus macaques"

### Figure S2

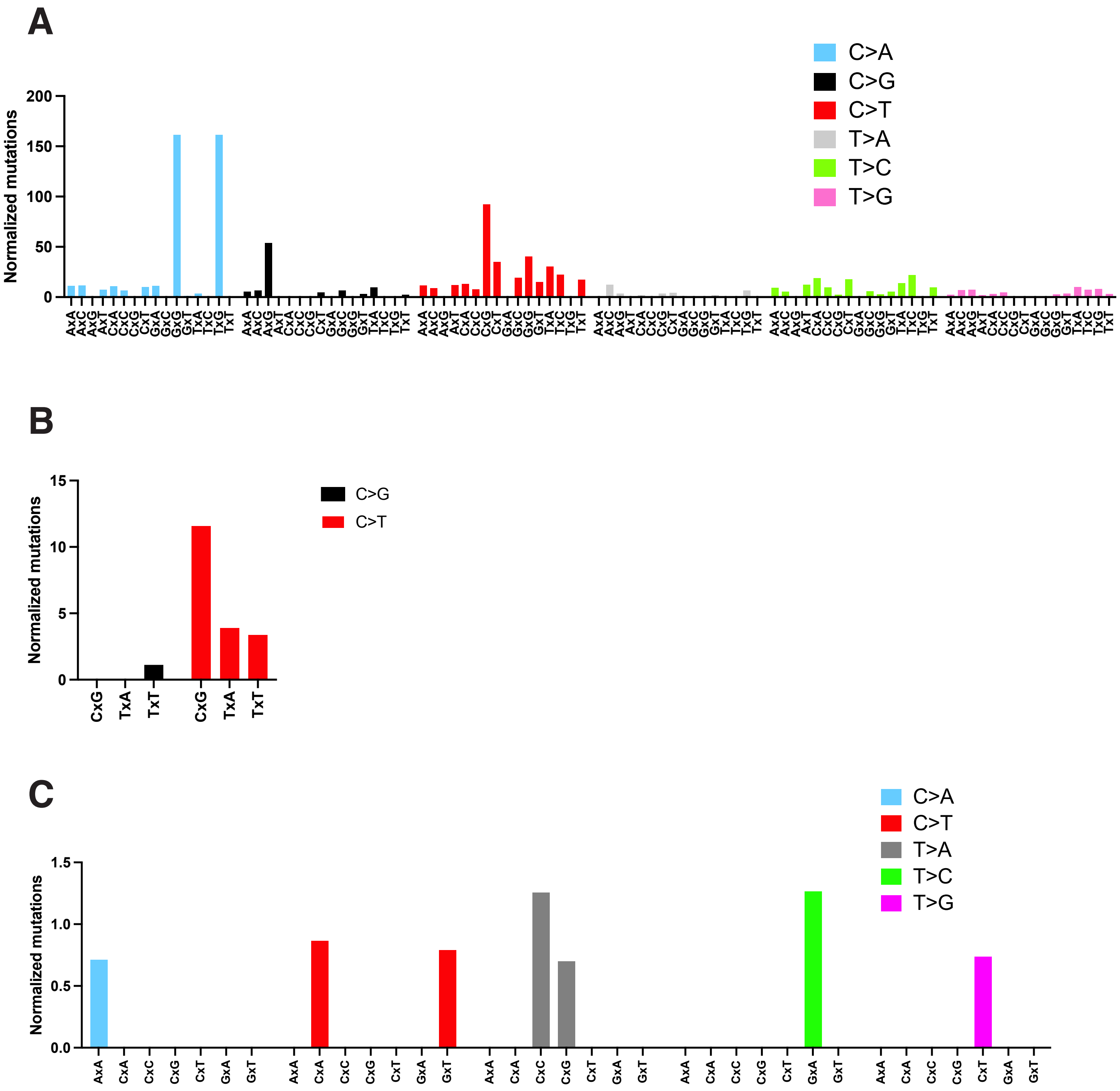

### Figure S3

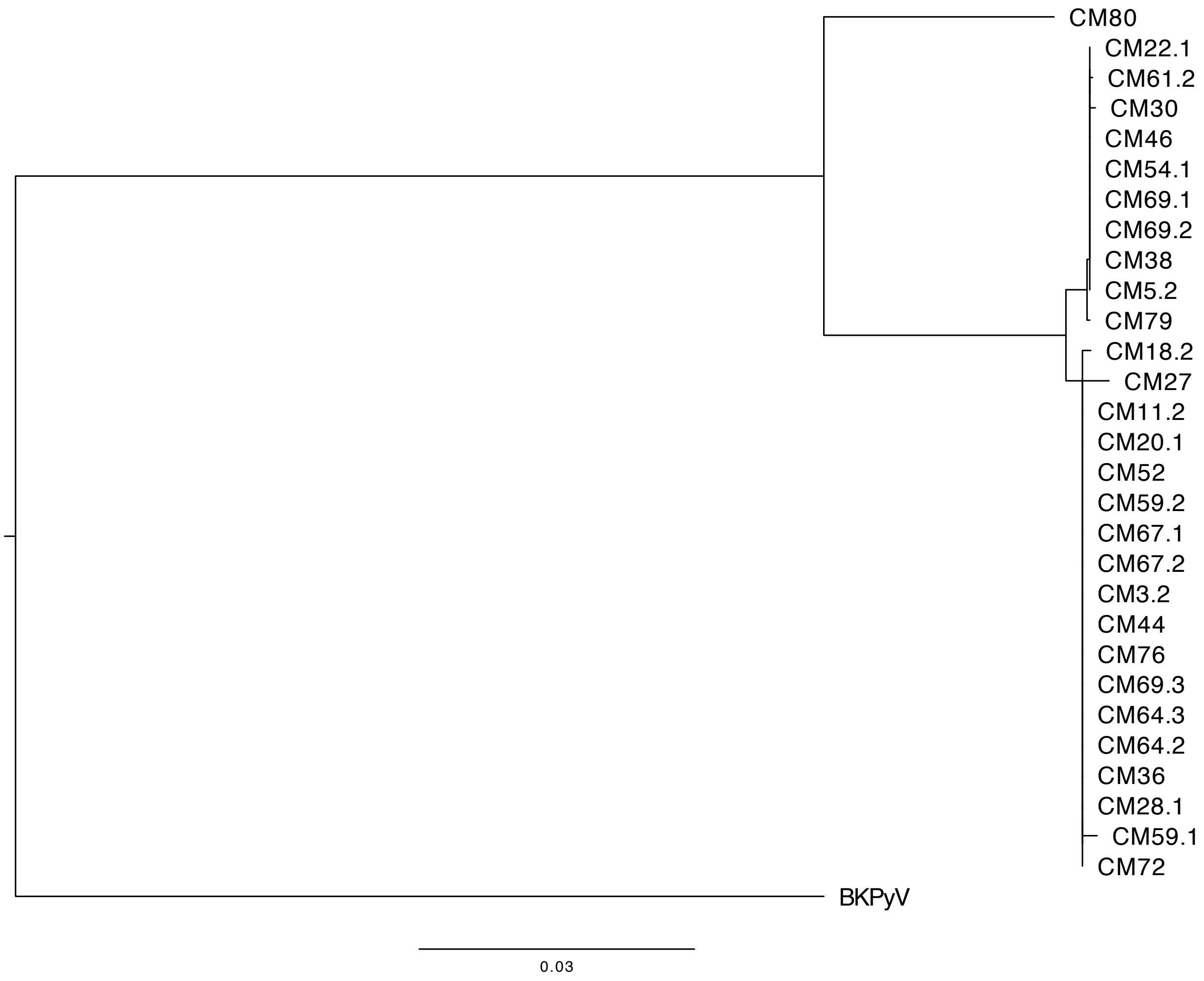
